# Understanding feeding competition under laboratory conditions: Rohu (*Labeo rohita)* versus Amazon sailfin catfish (*Pterygoplichthys* spp.)

**DOI:** 10.1101/2024.01.06.574454

**Authors:** Suman Mallick, Jitendra Kumar Sundaray, Ratna Ghosal

## Abstract

Competitive interactions between species is widely prevalent within the animal world. In this manuscript, we attempted to understand feeding competitions between the Amazon sailfin catfish, an invasive species introduced globally, and rohu, a keystone species native to several countries within southeast Asia. We used two different size classes of each species, large-size having total length (TL, from snout tip to caudal fin) of 15-20 cm and fingerling having TL <6 cm, and feeding duration was used as a proxy to understand competition. Our results demonstrated that feeding durations of large-size rohu were either similar or significantly (P<0.05) higher in presence of catfish when compared to trials in presence of conspecifics, indicating that large-size rohu is not a weak competitor. However, feeding durations of fingerling rohu was significantly (P<0.05) reduced in presence of both large-size and fingerling catfish, when compared to trials in presence of conspecifics. Moreover, fingerling rohu also displayed freeze (alarm) behavior in presence of the catfish. Interestingly, presence of rohu had no significant (P>0.05) impact on feeding durations of catfish. Overall, the study demonstrated that invasive catfish may behaviorally outcompete fingerling rohu, thus, threatening the sustenance of a species that is native to several freshwaters around the globe.

## Introduction

The competitive exclusion principle by Gause (1934), spurred a lot of interest among ecologists and ethologists to test the theory under varied laboratory and field conditions. Subsequent studies following Gause’s work demonstrated that competition between any two species may vary over temporal, for example, across developmental stages (Brenden and Murphy, 2004), across seasons (Borcherding et al., 2019), and spatial scales, for example, across habitats with different levels of anthropogenic disturbances (Carrete et al., 2010) and/or under conditions with varying levels of resources (Rudolf and McCrory, 2018). Fate of the interactions greatly depend upon the competitive ability of the interacting species (Birch, 1957; Bowers and Brown, 1982). For example, Gause showed *Paramoecium aurelia* outcompeted (in terms of population size) *P. caudatum* in combined cultures, however, the latter was the winner while competing with *P. bursaria* (Gause, 1934). Further, Gause’s experiments (1934) also demonstrated that due to competitive interactions for resources, for example, food and space, one of the interacting species may outcompete the other and eventually may lead the poorer competitor towards exclusion from the system or a given habitat. Such exclusions are of great concern if the species is a native one for a given habitat, and may lose out to a highly competitive exotic or alien species (Beisner et al., 2003), thus, impacting the stability of a given ecosystem (Jansen et al., 2014).

In this paper, we aim to understand competitive interactions in the context of feeding between two freshwater fish species, the Amazon sailfin catfish and rohu, that have been reported to co-occur in different countries across Asia. The Amazon sailfin catfish is native to the Amazon basin, and has been tagged as one of the invasive species in several countries, Mexico (Wakida-Kusunoki et al., 2007), Florida (Ludlow and Walsh, 1991), Texas (Edward, 2001), Malaysia (Saba et al., 2020), Iraq (Qasim and Jawad, 2022), Vietnam (Levin et al., 2008), Philippines (Jumawan et al., 2011), Sri Lanka (Herath et al., 2020), Italy (Piria et al., 2018), Bangladesh (Hossain et al., 2018), and the Indian freshwaters (Bijukumar et al., 2015; Rao and Sunchu, 2017; Hussan et al., 2019; Das et al., 2020; Seshagiri et al., 2021, Mallick et al., 2023). Amazon sailfin catfish is a benthic forager with hard armor and spines, belonging to the Loriicariidae family of catfish, which are considered to be aggressive and territorial (Hossain et al., 2018; Umar and Ramayani, 2021). Moreover, several species of catfish, for example, “*multiradiatus*”, “*pardalis*”, “*disjunctivus*”, belonging to the genus *Pterygoplichthys* have been introduced in different freshwater habitats across the globe (Singh and Lakra, 2011; Sumanasinghe and Amarasinghe, 2014; Tarkan et al., 2017). Due to high rate of hybridization among the different species of sailfin catfish, and due to difficulty in detecting hybrids using physical features, most studies refer to Amazon sailfin catfish using the genus name, *Pterygoplichthys* spp. (Jumawan et al., 2011).

Amazon sailfin catfish is a voracious algae eater and have been shown to have a huge dietary overlap with several native species in the introduced habitats (Gestring et al., 2010; Parvez et al., 2023). One such native species belonging to the Indian freshwaters is rohu *Labeo rohita* (Suresh et al., 2019). Rohu is a natural inhabitant of the Indian freshwaters (Das et al., 2013) and widely distributed in different rivers, for example, Cauvery, Narmada (Bhaumik et al, 2017), Tapti and Mahanadi (Mohanta et al., 2008). Rohu is considered as a keystone (Dwivedi et al., 2020) and an ecological indicator species (Mahmood et al., 2021), and is also economically valuable (Modeel et al., 2023). India is the second largest producer (9.59 million tonnes) of rohu (Seshagiri et al., 2022) contributing to 1.07% of national GDP along with an export income generation of 31,000 million, as reported in 2018-19 (Ngasotter et al., 2020). Using mesocosm-based approach, several studies have shown that Amazon sailfin catfish negatively impacted the growth of fingerling rohu [fingerling defined in terms of total length (TL, from snout to tip of caudal fin) of <15 cm, Munilkumar and Nandeesha, 2007], when both the species were reared together under similar experimental conditions (Parvez et al., 2023; Mallick et al., 2023). However, the catfish had no significant impact on different abiotic (water quality and soil C/N profiles) and biotic (zoo- and phytoplankton abundance) measures of the given experimental condition (Seshagiri et al., 2021; Parvez et al., 2023; Mallick et al., 2023). Based on the findings, the studies have speculated that the observed negative impact on rohu fingerlings can be mediated via feeding competitions with the catfish (Mallick et al., 2023), and thus, warrants further investigations. Thus, in this manuscript, we asked the following questions to understand feeding competition under laboratory conditions between Amazon sailfin catfish and rohu:

a. Characterizing feeding behavior of Amazon sailfin catfish: What kind of feeding behaviors are shown by Amazon sailfin catfish? To the best of our knowledge, there has been no studies on feeding behaviors of catfish, and we need to first define and characterize the behaviors in order to study competitive interactions.
b. Characterizing feeding behavior of Rohu: Do Rohu feed (in terms of feeding duration) on a given item provided in our laboratory conditions? Though rohu feeding behavior has been studied in detail (Mookerji and Rao, 1993; Rahman et al., 2008; Rahman, 2015), we wanted to find out if the species considerably fed on the items provided under our laboratory conditions to justify its inclusion in the competition trials.
c. Characterizing feeding interactions of Amazon sailfin catfish and rohu in presence of each other: Is there any impact on feeding behaviors (in terms of feeding duration) of rohu in presence of the Amazon sailfin catfish? If so, does it depend on the body size (TL, as a proxy for age, Raphael et al., 2016) of both species? Feeding duration of each species was used as a proxy to understand competition.

The purpose of the study is to understand feeding competitions between Amazon sailfin catfish and rohu to help understand any potential impact of the invasive catfish (Parvez et al., 2023; Wakida-Kusunoki et al., 2007) on the keystone and commerciallly valuable species, rohu (Dwivedi and Nautiyal, 2012; Dwivedi et al., 2020; Rabbane et al., 2022; Mallick et al., 2023). Such understanding may help in assessing the threats associated with several ecosystems across the globe, where the native rohu now co-occur with the invasive catfish (Wakida-Kusunoki et al., 2007; Saba et al., 2020; Seshagiri et al., 2021; Parvez et al., 2023).

## Materials and Methods

### Study animals

Rohu and catfish were broadly divided into two size classes based on TL: large (TL=15-20 cm) and small or fingerlings (TL<6 cm). In fishes, length is used as a proxy for age, where younger ones have shorter length when compared to older fish (Raphael et al., 2016). Body weights were kept similar for both the species proportional to their respective size classes. Amazon sailfin catfish of the large-size class had an average length and weight of 17.76±0.31 cm (values here and throughout the manuscript presented as Mean±Standard Error [SE]) and 18.1±0.66 g, respectively. Rohu of the large-size class had an average length and weight of 18.9±0.20 cm and 18.04±5.09 g, respectively. Catfish fingerlings had an average length and weight of 5.85±0.10 cm and 2.18±0.08 g, respectively. Similarly, rohu fingerlings had an average length and weight of 4±0.12 cm and 2.06±0.05 g, respectively. Amazon sailfin catfish were procured from the local pet store whereas rohu were obtained from commercial aquaculture farms. Amazon sailfin catfish and rohu were stocked separately in 70 L glass tanks (53 x 37 x 37 cm) and were fed daily with algal wafer having a mixture of algae, shrimp meal, and fish oil (Red China, Guangzhou, China) for four months before the start of the experimental trials. Both the species had no prior exposure to each other before the experimental trials. Fishes were housed under laboratory conditions with a photoperiod of 14:10 light:dark cycle at 24-26**°**C. There were no observed mortality of the fishes during the entire experimental duration (6 months).

### Experimental design

Experiments were conducted under low light and low sound conditions. All the trials were conducted for a period of three hours and were recorded using a handycam (VIXIA HF R800, Canon, China). All the individuals were starved for 48 hours prior to the experiment. Trials were conducted during the day (09:00 to 13:00), and different behaviors in the context of feeding were identified, scored and defined during the playback of the recorded videos. Trials were initiated with the introduction of food bags that were made by wrapping 100 mg of algal wafer (Red China, Guangzhou, China) in a fine mesh and stitched on all the sides. From here onwards, we refer to algal wafer as ‘home food’ that was used during housing. Our pilot trials with catfish and rohu (N=4 trials for each species; each for 3 h) indicated that both the species were able to access food via the mesh, and the bags were found empty (food was eaten completely) after the said duration of the trials. Based on the results of the pilot trials, the main experimental scheme was decided as described in Fig. 1. All the trials for the main experiment (Trial type I, II, and III, as described below) were conducted in 70 L glass tanks with two individuals (either of single or mixed species depending on the type of the trials, Trial I, II and III, as mentioned below), and three such tanks were set up as biological replicates. Three trials were conducted on each tank (N=9 trials) after a gap of 48 h to provide a rest period to the experimental fishes before reuse.

**Fig. 1.**
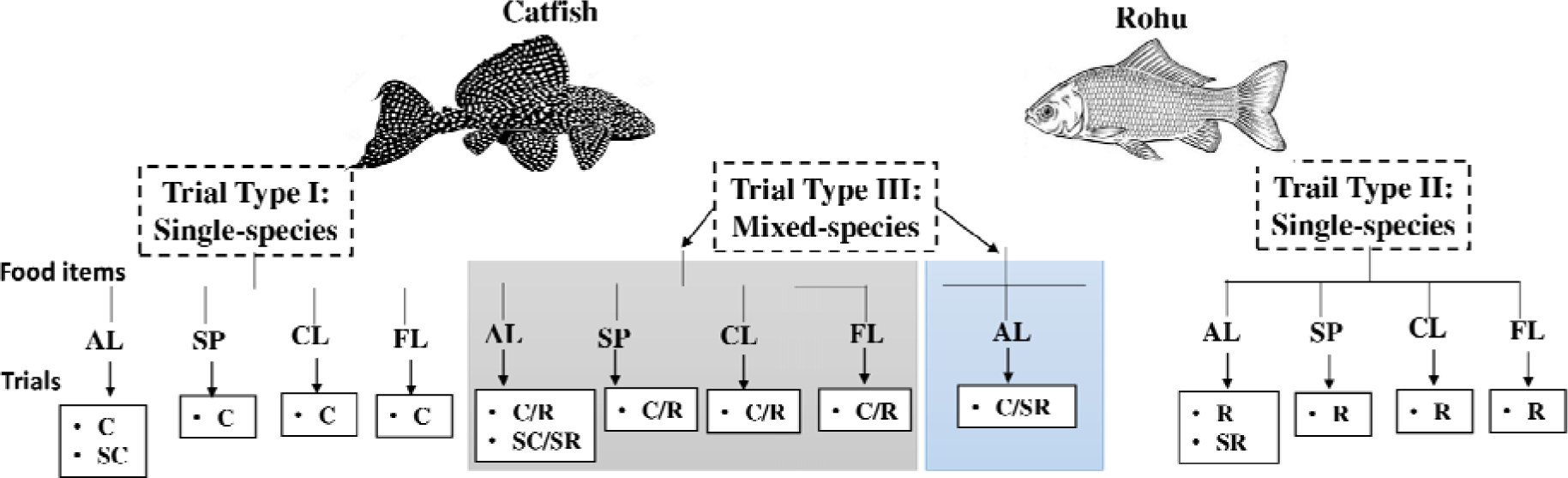
Schematic representation of the experimental design for assessing feeding competition between Amazon sailfin catfish and rohu. Experimental trials were classified as single-species (Trial Type I and II) and mixed-species (Trial Type III), while the latter was further divided into size-matched () and size-unmatched () classes. Food items used during the trials were algal wafer (AL), spirulina (SP), chlorella (CL), and flake (FL). All large-size, single species trials were denoted as C and R, all fingerling trials as SC and SR, for catfish and rohu respectively. Mixed species trials for size-matched category were C/R for large-size class and SC/SR for fingerlings, and for size-unmatched having large-size catfish and fingerling rohu, was C/SR.

### Trial type I: Single-species trials to characterize feeding behavior of the Amazon sailfin catfish

Feeding trials for large-size catfish are abbreviated as ‘C’ from now onwards, and that of fingerling catfish trials are abbreviated as ‘SC’. A similar protocol, as mentioned above was followed for C and SC trials. C trials were conducted with four different food items, ‘home food, spirulina (composed of blue-green algae, Parry Limited, Tamil Nadu, India), chlorella (composed of single celled green algae, Parry Limited, Tamil Nadu, India), and flake (composed of fish meal, Taiyo Feed Mill Pvt. Ltd., Tamil Nadu, India) added separately (N=9 trials for each food item) to the tanks. Since catfish is known to be an algal eater (Gestring et al., 2010; Parvez et al., 2023), most of the selected food items were algae-based. Durations of different behaviors including feeding were calculated for each food item during the playback of the recorded video of the trials. However, SC trials (N=9 trials) were conducted only using the ‘home food’.

### Trial type II: Single-species trials to characterize feeding behavior of rohu

Feeding trials for large-size rohu are abbreviated as ‘R’ from now onwards, and that of fingerling rohu trials are abbreviated as ‘SR’. Experimental conditions were the same as described above for trial type I. Apart from the ‘home food’, R trials were conducted with the same three food items, spirulina, chlorella and flake, as described above for C trials, and previous studies have shown these items to be preferred by rohu (Rahman, 2015). The food items were added separately to the rohu tanks (N=9 trials for each food item, spirulina, chlorella, flake and ‘home food’). SR trials (N=9 trials) were conducted with only the ‘home food’. Durations of different behaviors including feeding were calculated for each food item during the playback of the recorded video of the trials.

### Trial type III: Mixed-species trials to characterize interspecific interactions between the Amazon sailfin catfish and rohu

The mixed-species trials were categorized into size-matched and size-unmatched setups. Size-matched trials were conducted using individuals of equal body size (TL) and matching body weights for both the species, rohu and Amazon sailfin catfish. Size matched trials were of two types, large-size catfish with large-size rohu, abbreviated as C/R from now on; and fingerling catfish with rohu fingerling, abbreviated as SC/SR from now on. For C/R trials, we used one catfish and one rohu, both species having a matching length of ∼17-18 cm and weight of ∼17-18 g for each individual. However, for SC/SR, we used four individuals of each species, both species having matching length of 4-5 cm and weight of 1-2 g for each individual. We used four fingerlings of each species for SC/SR trials as fingerling (young) fish usually show reduced feeding when occurring under solitary conditions (Boujard, 1995). C/R trials were conducted for all the four food items, algal wafer, spirulina, chlorella and flake (N=9 trials for each food item), added separately under similar experimental conditions described above. SC/SR trials (N=9) were conducted only on the ‘home food’. For both C/R and SC/SR, durations of different behaviors including feeding were calculated for each food item during the playback of the recorded video of the trials.

With regards to the size-unmatched category, trials were conducted by including large-size catfish with rohu fingerlings, abbreviated as C/SR trials from now on. For size-unmatched trials (C/SR), biomass (weight) matching (de Lourdes Ruiz-Gomez et al., 2008) was taken into consideration by keeping 8 fingerling rohu (having a length of ∼4-5 cm and a weight of ∼2-2.5 gm for each individual) with one large-size catfish (length of ∼17-18 cm and weight of ∼17-18 g). C/SR trials (N=9) were conducted only on the ‘home food’.

For all mixed-species trials, durations of different behaviors including feeding were calculated for each food item during the playback of the recorded video of the trials, from the perspective of both catfish and rohu species.

### Statistical analysis

All the behavioral durations are expressed as Mean±Standard Error (SE) per individual per trial for both mixed and single species’ trials. Behavioral data (durations) was tested for normality by conducting Shapiro-Wilk test, and F-test was performed to check for equal or unequal variances. Following normality assumptions (raw data or after log or log x+1 transformations), Student’s t-test was applied to data to compare between two groups of treatment that had equal variances. Comparison between two groups having unequal variances was conducted using Welch’s t-test. Mann-Whitney U test was performed to compare between two groups of treatment that followed non-normal distribution. Accordingly, One-way ANOVA or Kruskal-Wallis test was performed for comparisons across more than two groups. The level of significance was kept at P≤0.05. All the statistical analyses were done on the R program, version 4.2.3. For all the mixed-species (rohu and catfish) trials (both size-matched and -unmatched categories), durations of different behaviors displayed by a species, particularly feeding durations, were compared and contrasted with the durations in their respective single species’ (in presence of the conspecific) trials in order to understand competition between the two species.

## Results

### Trial type I: Single species trials to characterize feeding behavior of the Amazon sailfin catfish

Three behaviors, feeding, push and aggressive attack, were characterized, defined and measured in the C and SC trials. A detailed catalog of different behaviors displayed by the catfish in the context of feeding has been listed in Table 1 along with the duration of each behavior. Feeding and push behaviors were displayed by both large-size and fingerling catfish whereas aggressive attack was displayed only by large-size catfish.

**Table 1:** Description of different behaviors of catfish and rohu in the context of feeding using the ‘home’ food, algal wafer.

| Behaviors | Description | Duration (s) or Frequency*<br>(Mean±SE per individual per trial) |  |  |  |
| --- | --- | --- | --- | --- | --- |
|  |  | Large-size<br>catfish<br>in<br>C trials | Fingerling<br>catfish in<br>SC trials | Large-size<br>rohu in<br>R trials | Fingerling<br>rohu in<br>SR trials |
| **Feeding** | Placing suckers (for catfish) or mouth (for rohu) over a food item accompanied by rapid buccal movements. | 1185.61<br>±<br>262.97 | 1654.13±<br>131.29 | 448.75±<br>95.33 | 525.96±<br>77.82 |
| **Push** | Striking the other individual with the snout or dorsal fin to drive away from food. | 62.94±20<br>.89 | 76.66±20.5<br>1 | 190.58±<br>127.8 | Not<br>observed |
| **Aggressive**<br>**attack** | Rapid movement with erected dorsal fin towards the other individual. | 15.55±<br>7.10 | Not<br>observed | Not<br>observed | Not<br>observed |
| **Freeze** | Cessation of movement and swimming activity for ≤10 sec. | Not<br>observed | Not<br>observed | Not<br>observed | 32.18±1.85 |

\*All behaviors are expressed in durations (duration/individual/trial) except freeze behavior, which is calculated as frequency (number of times/individual/trial).

Feeding durations of large-size catfish (in C trials) were similar across the four tested food items, ‘home food’ (1185.61±262.97 s), flake (771.05±109.43 s), chlorella (627.72±135.99 s), spirulina (648.77±59.40 s), with no significant differences (One-way ANOVA; F=2.6; P=0.06) across the trials (Supplementary Fig. 1a). For comparison purpose, from here onwards, we pooled the durations of aggressive attacks and push behavior to represent aggressive interactions. Duration of aggressive interactions of large-size catfish for the four tested food items were 78.5±23.15 s for home food, 165.22±43.85 s for flake, 65.94±28.81 s for chlorella, and 197.22±50.42 s for spirulina (Supplementary Fig. 2a). The duration of aggressive interactions of large-size catfish differed significantly (One-Way ANOVA; F=3.61; P=0.02) across the four tested food items (Supplementary Fig. 2a), and post-hoc Tukey test showed significant differences (P<0.05) between chlorella and spirulina.

The feeding duration of fingerling catfish (in SC trials) was 1654.13±131.29 s for ‘home food’ and duration of push behavior was 76.66±20.51 s. Since fingerling catfish did not show any aggressive attacks, we included push behavior only to represent aggressive interactions of fingerlings for further comparisons.

### Trial type II: Single species trials to characterize feeding behavior of rohu

Three behaviors, feeding, push and freeze were characterized, defined and measured in the R and SR trials. A detailed catalog of different behaviors displayed by rohu including feeding has been listed in Table 1 along with the duration or frequency of each behavior. Push behavior was displayed by large-size rohu only, whereas freeze behavior was only displayed by fingerling rohu (Table 1).

Feeding durations for large-size rohu was 448.75±95.33 s for ‘home food’, 496.5±130.67 s for flake, 1437.33±349.84 s for chlorella, and 1863.83±439.10 s for spirulina. Feeding durations of large-size rohu were significantly different (One-way ANOVA; F=5.77; P=0.005) across the tested food items (Supplementary Fig. 1b), and post-hoc Tukey test showed significant differences (P<0.05) in feeding durations between algal wafer and spirulina, and flake and spirulina (Supplementary Fig. 1b). No significant differences (P>0.05) were obtained in pairwise comparisons (post-hoc Tukey test) for the rest of the food items. However, there were no significant differences in the duration of push behavior (represented as aggressive interactions) across the tested food items (Kruskal wallis test; ^2^=0.65; P=0.88) (Supplementary Fig. 2b). Duration of aggressive interactions for large-size rohu was 190.58±127.80 s when presented with algal wafer, 32.83±28.57 s with flake food, 44.25±29.23 s with chlorella, and 14.75±6.81 s with spirulina.

Feeding duration of fingerling rohu (in SR trials) was 525.9±77.82 s for ‘home food’ and no push behavior was displayed by fingerling rohu. Freeze behavior was displayed by fingerling rohu having a frequency of 32.18±1.85/individual/trial, however, freeze was observed only during the mixed-species trials, and the details are mentioned below.

### Trial type III: Mixed-species trials to characterize interspecific interactions between Amazon sailfin catfish and rohu

In C/R trials for four different food items, the average feeding duration of large-size rohu, in absolute terms, was either higher or equal when compared to durations in (single-species) R trials (Fig. 2a-d). Feeding durations of rohu in C/R trials were 1696±336.03 s for ‘home food’, 1801.16±460.83 s for spirulina, 1678.5±267.88 s for chlorella and 1275.66±293.14 s for flake. Significantly higher feeding durations were observed in large-size rohu during C/R trials for algal wafer (Welch’s t-test; t=3.57; P=0.01) (Fig. 2a) and flake (Student’s t-test; t=2.42; P=0.03) (Fig. 2d) when compared to R trials for the given food items. No significant differences were obtained in feeding durations of large-size rohu between C/R and R trials for spirulina (Student’s t-test; t=-0.09; P=0.92) (Fig. 2b) and chlorella (Student’s t-test; t=0.54; P=0.59) (Fig. 2c) items. Similar pattern was observed for large-size catfish in C/R trials. In absolute terms, large-size catfish, on an average, showed higher durations of feeding in mixed-species C/R trials when compared to single-species C trials. The feeding durations of large-size catfish in C/R trials were 2677.66±697.51 s for home food, 1318±460.83 s for spirulina, 2581± 271.50 s for chlorella, 2692.5±852.61 s for flake. Significant differences were observed in feeding duration of large-size catfish when compared between C/R and C trials for three of the food items, algal wafer (Student’s t-test; t=-2.30; P=0.03) (Fig. 2a), spirulina (Mann-Whitney U test, W=0, P=0.0003) (Fig. 2b) and chlorella (Student’s t-test; t=-7.09; P=8.061e-06) (Fig. 2c). No significant differences were obtained for feeding durations of large-size catfish between C/R and C trials for flake food (Welch t-test; t=-2.23; P=0.07) (Fig. 2d).

**Fig. 2.**
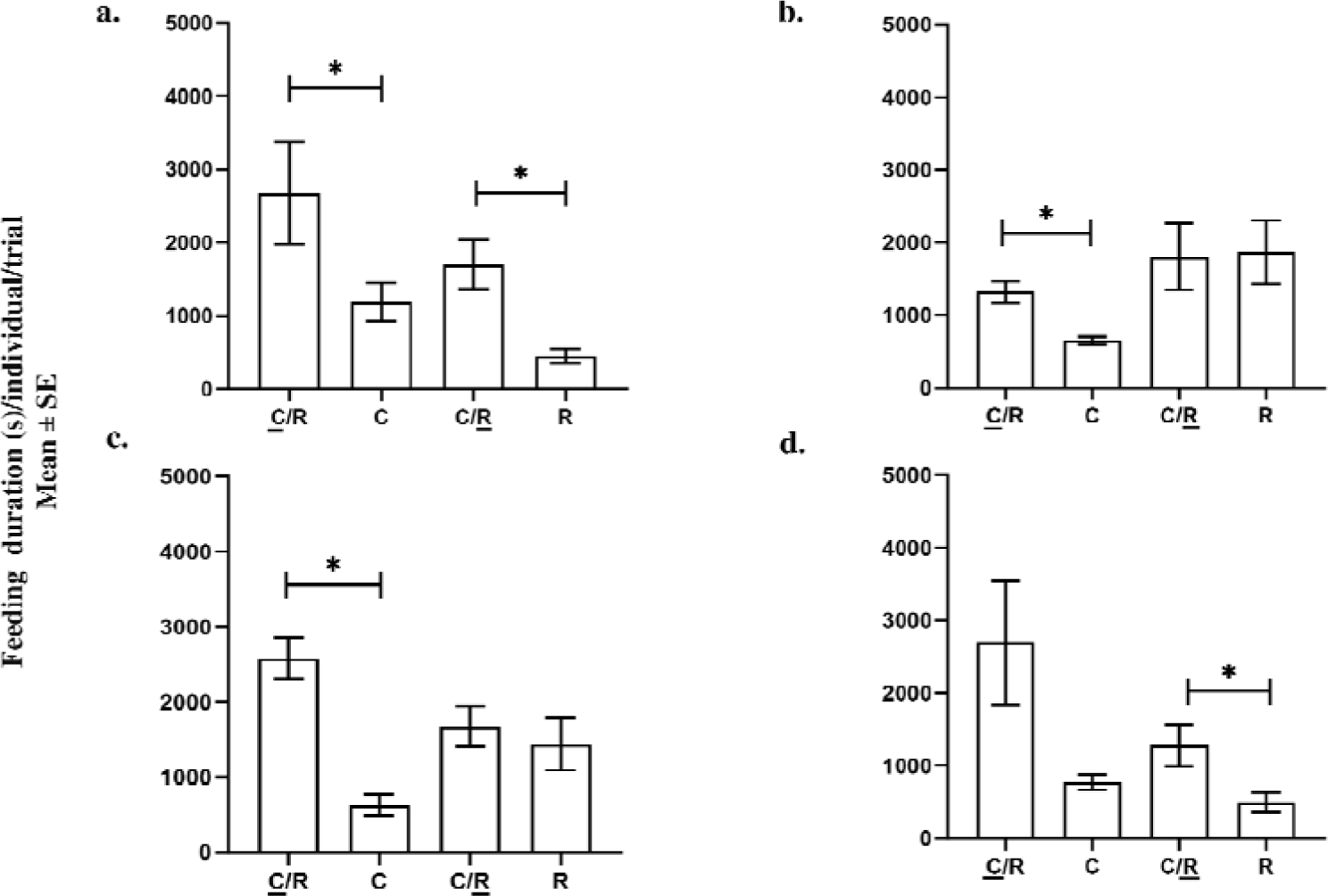
Feeding durations (Mean±SE per individual per trial) of large-size Amazon sailfin catfish in C trials (N=9 trials for each food item), large-size rohu in R trials (N=9 trials for each food item), and large sizes of both Amazon sailfin catfish and rohu in size-matched, mixed-species C/R trials (N=9 trials for each food item) for different food items, panel a: algal wafer; panel b: spirulina; panel c: chlorella; and panel d: flake food. The underlined <u>C</u> and <u>R</u> represent feeding durations of large-size catfish and large-size rohu in C/R trials, respectively. *indicates significant differences (P≤0.05). ‘+’ represents the mean value per individual per trial.

The duration of aggressive interactions of large-size catfish in C/R trials for four tested food items (Fig. 3) were 150.66±27.86 s for home food, 137.16±22.87 s for spirulina, 62±13.98 s for chlorella, 127.83±31.75 s for flake. There were no significant differences in the duration of aggressive interactions of large-size catfish between C and C/R trials for spirulina (Welch’s t test, t=0.21; P=0.83) (Fig. 3b), chlorella (Welch’s t test, t=-1.36; P=0.20) (Fig. 3c) and flake (Welch’s t test, t=0.69, P=0.50) (Fig. 3d). However, there was a significant difference (Student’s t-test; t=-3.89; P=0.003) in the duration of aggressive interactions of catfish between C and C/R trials for algal wafer (Fig. 3a). The duration of aggressive interactions of large-size rohu in C/R trials were 60.16±22.73 s for home food, 26.5±13.66 s for spirulina, 4±2.04 s for chlorella, 30.5±13.09 s for flake. There were no significant differences (Student’s t-test for algal wafer [t=0.82, P=0.42] and spirulina [t=0.76, P=0.45]; Mann-whitney U test for chlorella [W=20, P= 0.79] and flake [W=26, P=0.22]) in the duration of aggressive interactions of large-size rohu between R and C/R trials for all the tested food items (Fig. 3a-d).

**Fig. 3.**
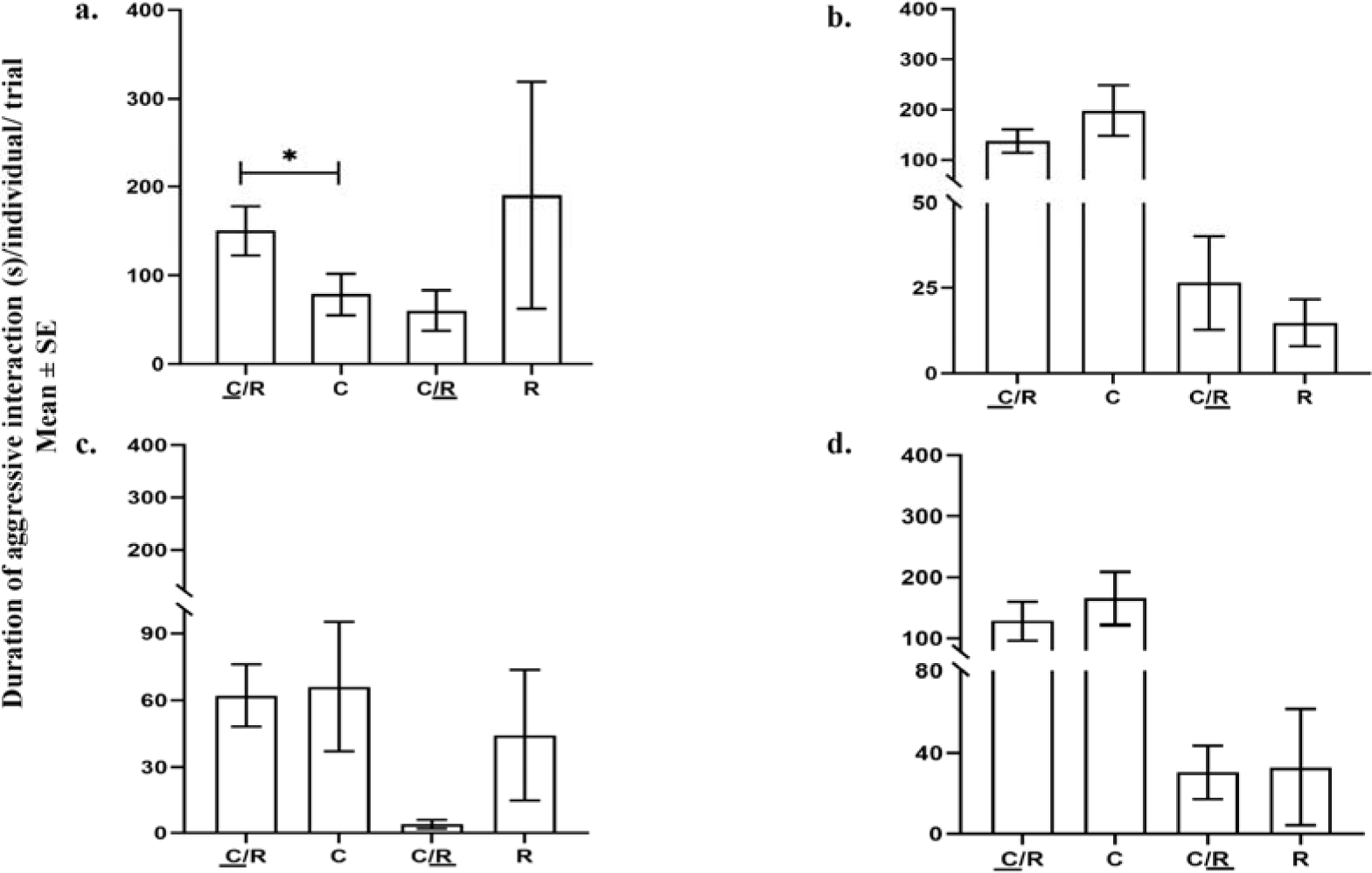
Duration of aggressive interactions (Mean±SE per individual per trial) of large-size Amazon sailfin catfish in C trials (N=9 trials for each food item), large-size rohu in R trials (N=9 trials for each food item) and large-size of both catfish and rohu in size-matched, mixed-species C/R trials (N=9 trials for each food item) for different food items, panel a: algal wafer; panel b: spirulina; panel c: chlorella; and panel d: flake food. The underlined <u>C</u> and <u>R</u> represent durations of aggressive interactions of large-size catfish and large-size rohu in C/R trials, respectively. *indicates significant differences (P≤0.05). ‘+’ represents the mean value per individual per trial.

In SC/SR trials, fingerling rohu showed significantly (Welch’s t-test; t=6.70; P=0.001) reduced feeding duration (85.34±33.28 s) in presence of fingerling catfish when compared to single-species SR trials (Fig. 4). In contrast, fingerling catfish showed no significant (Student’s t-test; t=-1.99; P=0.07) differences in feeding duration (1979.20±148.49 s) in presence of fingerling rohu in SC/SR trials when compared to single-species SC trials (Fig. 4). In size-unmatched mixed-species C/SR trials, feeding duration of fingerling rohu (192.02±36.16 s) significantly (Welch’s t-test; t=-3.89; P=0.003) reduced when compared to SR trials (Fig. 5). However, there were no significant (Welch’s t-test; t=-1.86; P=0.11) differences in feeding duration (3670.5±1308.18 s) of large-size catfish between C/SR and C trials (Fig. 5). In fact, in absolute terms, the average feeding duration of large-size catfish was higher in C/SR when compared to C trials (Fig. 5).

**Fig. 4.**
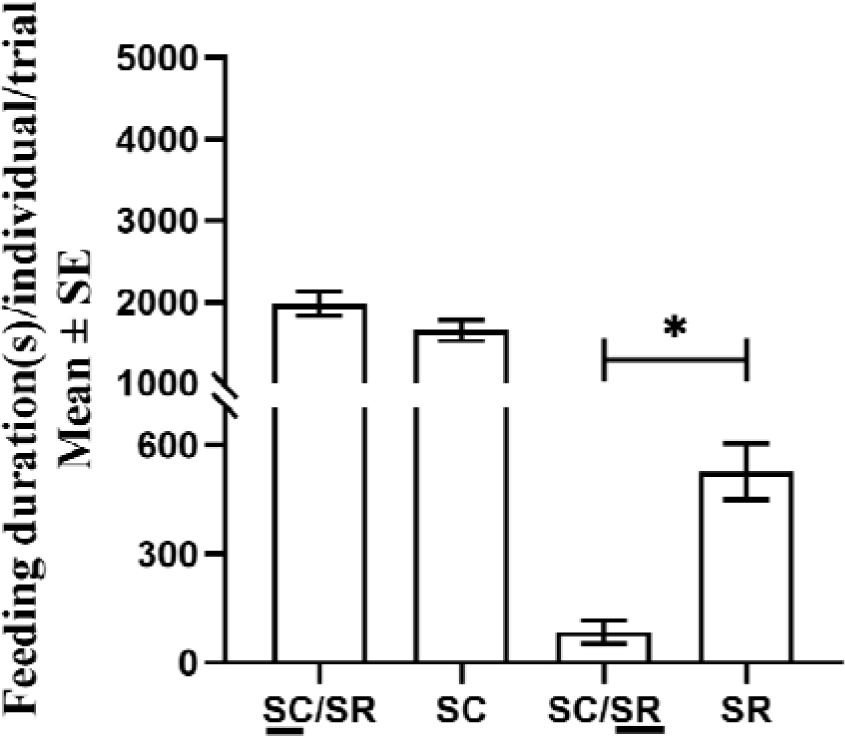
Feeding durations (Mean±SE per individual per trial) of fingerling catfish in SC trial (N=9 trials), fingerling rohu in SR trials (N=9 trials) and fingerlings of both catfish and rohu in size-matched, mixed-species SC/SR trials (N=9 trials). The underlined <u>SC</u> and <u>SR</u> represent feeding durations of fingerling catfish and fingerling rohu in SC/SR trials, respectively. *indicates significant differences (P≤0.05). ‘+’ represents the mean value per individual per trial.

**Fig. 5.**
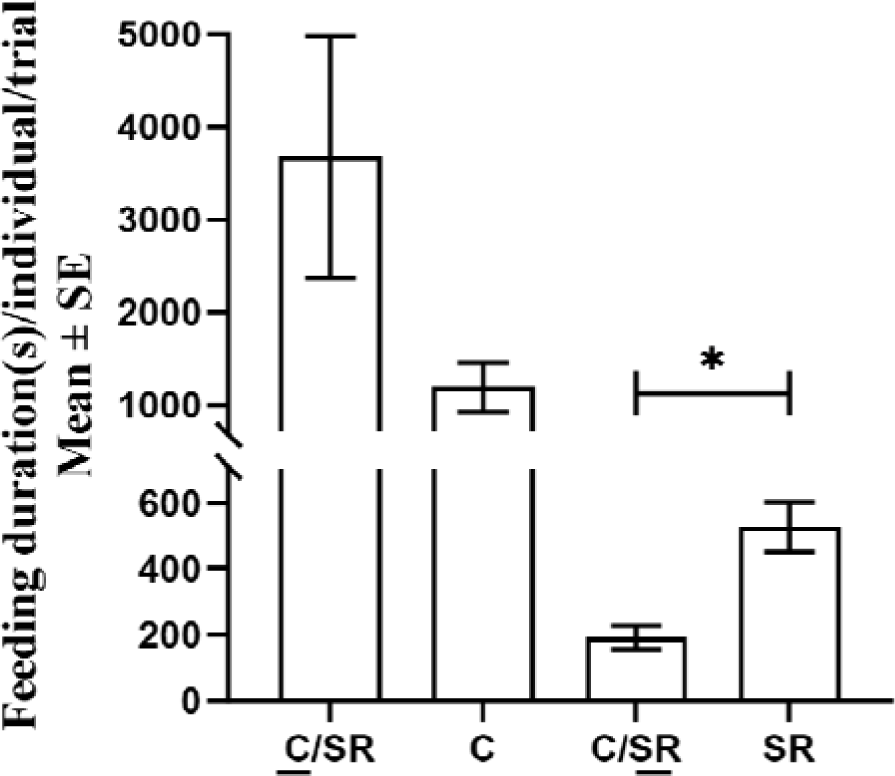
Feeding durations (Mean±SE per individual per trial, N=9 trials) of large-size catfish and fingerling rohu in size-unmatched, mixed-species C/SR trials. The underlined <u>C</u> and <u>SR</u> represent the feeding durations of large-size catfish and fingerling rohu in C/SR trials, respectively. Feeding durations of large-size catfish and fingerling rohu in C and SR trials are repeated from Fig. 2 and Fig. 4, respectively for comparison purposes only. ‘+’ represents the mean value per individual per trial.

For fingerling catfish, duration of aggressive interactions (23.33±4.69 s) reduced significantly (Welch’s t-test; t=2.76; P=0.03) (Fig. 6) in SC/SR trials when compared to SC trials. Similarly, for size unmatched C/SR trials, duration of aggressive interactions of large-size catfish (16±4.65 s) reduced significantly (Welch’s t-test; t=-3.89; P= 0.005) when compared to C trials (Fig. 6). Since fingerling rohu did not show any aggressive behavior, no comparisons could be made.

**Fig. 6.**
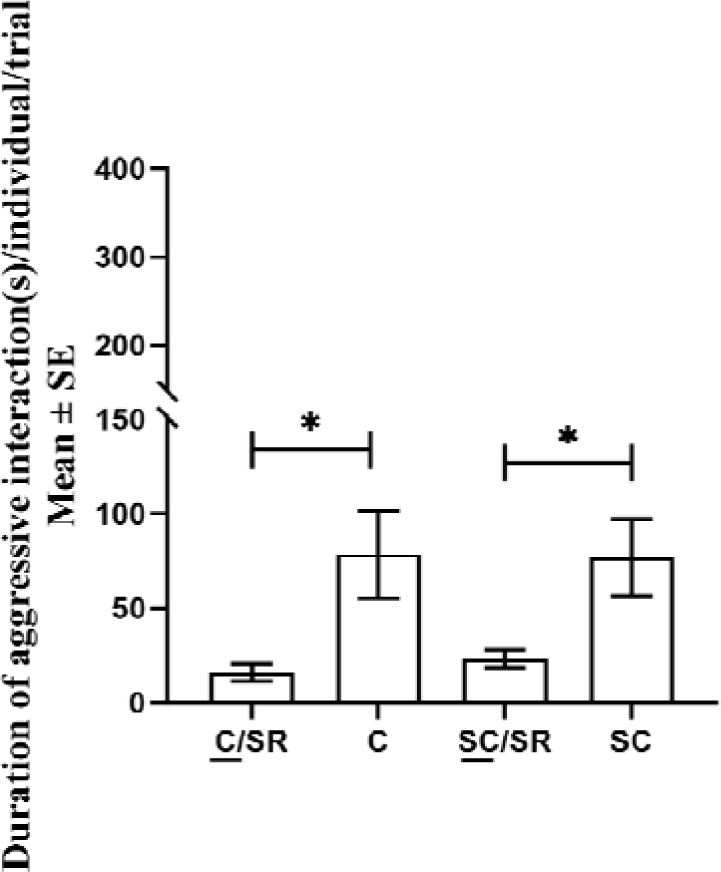
Duration of aggressive interactions (Mean±SE per individual per trial) of fingerling catfish in single-species SC trials (N=9 trials), fingerling catfish in size-matched, mixed-species SC/SR trials (N=9), and large-size catfish in size-unmatched, mixed species C/SR trials (N=9 trials). The underlined <u>C</u> and <u>SC</u> represent durations of aggressive interactions of large-size and fingerling catfish in C/SR and SC/SR trials, respectively. Duration of aggressive interactions of large-size catfish in C trials is repeated from Fig. 3 for comparison purposes only. *indicates significant differences (P≤0.05). ‘+’ represents the mean value per individual per trial.

Freeze behavior was displayed only by fingerling rohu both in C/SR and SC/SR trials (Fig. 7) The frequency of freeze behavior in C/SR trials was 52.12±3.71 and in SC/SR trials was 12.18±2.77. A significant (Student’s t-test; t=8.60; P=6.155e-06) difference was observed in freeze behavior between C/SR and SC/SR trials, wherein fingerling rohu showed increased freeze behavior in presence of large-size catfish when compared to fingerling catfish.

**Fig. 7.**
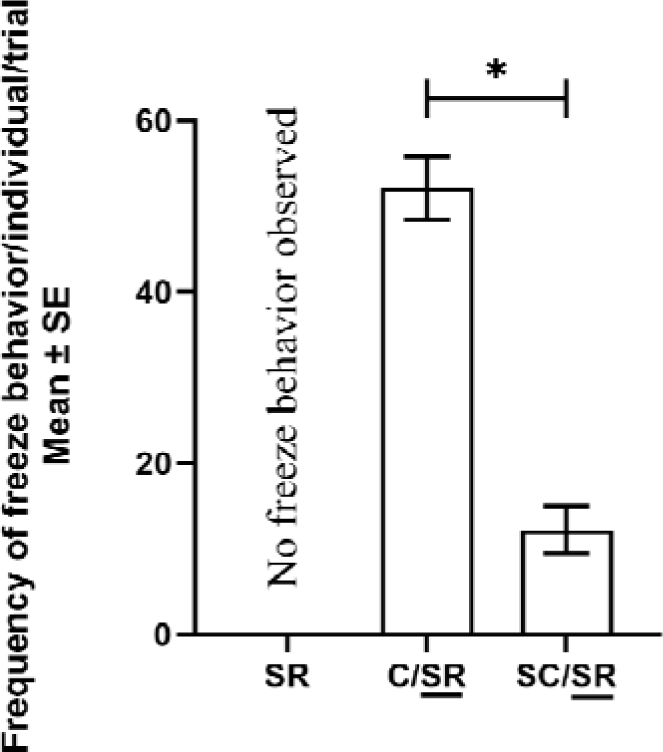
Frequency of freeze behavior (Mean±SE per individual per trial, N=9 trials) of fingerling rohu (represented as underlined <u>SR)</u> in size-matched and size-unmatched, mixed-species SC/SR and C/SR trials. *indicates significant differences (P≤0.05). ‘+’ represents the mean value per individual per trial.

## Discussion

The study, for the first time, described feeding behavior of Amazon sailfin catfish under laboratory conditions showing that the feeding interactions in catfish is frequently associated with aggression. Our results demonstrated that rohu fingerlings were weak competitors in presence of catfish, and showed a significant reduction in feeding duration when compared to durations in presence of the conspecifics. This reduction in feeding durations of fingerling rohu was independent of the size of the catfish, and occurred in presence of both fingerling and large-size catfish. Further, rohu fingerlings also showed freeze behavior in presence of the catfish, which could be a sort of alarm response towards a hard-armor, aggressive species. Contrastingly, large-size rohu and large-size catfish had comparable or even higher feeding durations in presence of each other when compared to durations in presence of their conspecifics for a given food item. Thus, both the species of the large-size category appeared to be equal in terms of their competitive abilities in the context of feeding. However, whether the dynamics of feeding competition between the two species change in relation to varying levels of resources (from low to high food availability), and across temporal (breeding versus non-breeding seasons) and spatial (different habitats) scales warrants further investigations (Glova, 1986; Balshine et al., 2005; Grabowska et al., 2016).

Our study demonstrated that feeding behavior of catfish is mostly associated with aggression (Hossain et al., 2018; Umar and Ramayani, 2021), which is characteristic of both fingerling and large-size classes, and is displayed towards conspecifics and hetero species, as well. Though there was a reduction in aggressive behavior of catfish towards rohu fingerlings when compared to aggression shown towards conspecifics (in C and SC trials) or large-size rohu (in C/R trials). This was not surprising as rohu fingerling was a weak competitor (in terms of reduced feeding duration) in presence of catfish, and thus, received less aggression from the catfish (Franco and Arce, 2022). Interestingly, in C/R trials, both species of large-size classes showed comparable feeding durations, and displayed similar or even higher levels of aggression towards each other (Miyai et al. 2011; McCarthy et al., 1999) when compared to the respective conspecific (C or R) trials. Thus, large-size rohu was not a weak competitior in presence of the large-size catfish and this pattern was consistent, in terms of both feeding and aggressive interactions, across different food items. Similar results, where different fish species competed equally in a given habitat and had no significant impact on each other, were demonstrated in several studes by Paszkowski (1985), Savino and Kolar (1996) and Fullerton et al., (2000). However, contrasting results were also demonstrated by Mills et al., (2004) showing that both large- and small-size classes of least chub (*Iotichthys phlegethontis*), a native fish to Bonneville basin of Utah, were behaviorally submissive in presence of mosquitofish (*Gambusia affinis*), an invasive fish within the same habitat. Thus, feeding competitions between species may vary depending on the competitive ability of the interacting partner, and needs thorough investigations under different environmental and ecological conditions.

The results for large-size rohu are in contrast to the findings observed for the fingerling rohu, which was outcompeted, in terms of feeding durations, in presence of both size-matched and large-size catfish. Similar results, where young individuals are poorer competitors than older ones, have been obtained by Winemiller and Taylor (1987), Persson and Greenberg (1990), and Byström and García-Berthou (1999) using different species of fishes, and such differences in competitive abilities of young verus old individuals could be due to experience or in relation to health or physiological characterisitics of a particular developmental stage (Nol and Smith, 1987; Šárová et al., 2013). Significant negative impact of the catfish on feeding durations of the fingerling rohu clearly draws parallels between the current work under laboratory conditions, and previous field-based studies (Mallick et al., 2023; Parvez et al., 2023), which also showed that growth of only fingerling rohu (and not larger than fingerling size class) got severely impaired in presence of the Amazon sailfin catfish within a given mesocosm setup. Similar impact of an exotic species, exclusively targeting young-age individuals of the native fish species, has been widely documented in several studies by Mills et al. (2004) and Bašić et al. (2019). Additionally, both the fingerling and large-size catfish were significantly impactful in terms of reducing feeding durations of rohu fingerlings. This is highly concerning in the field of invasion ecology and fisheries sciences, as Amazon sailfin catfish species has now spread in several Asian (Bijukumar et al., 2015; Parvez et al., 2023) and Southeast Asian countries (Jumawan et al., 2011; Chaichana et al., 2011), where Rohu is an important economically valuable, native species (Rasal and Sundaray, 2020; Modeel et al., 2023).

Poorer competitive ability of the fingerling rohu was further supported by the display of freeze behavior by fingerlings in presence of the catfish. Fingerling rohu displayed higher frequency of freeze behavior in presence of large-size catfish when compared to fingerling catfish, and such contrast might be obvious due to sheer size differences (Katsuya 2002; Templeton et al., 2005). Freeze behavior has been widely documented in several fish species, mostly displayed in the context of alarm or anti-predatory behavior (Brown et al., 1999; Bass and Garlai, 2008; Näslund et al., 2016). The results of this study are in agreement with the study on zebrafish (*Danio rerio*) (Bass and Garlai 2008), where the authors documented a significant difference in freeze behavior of zebrafish between test, in presence of an exotic swordtail fish (*Xiphophorus helleri*), and control, in presence of conspecifics, groups. Our study clearly demonstrates that the presence of the exotic catfish not only impedes the feeding duration of the fingerling rohu but also has a significant impact on the behavioral expressions of the rohu, which in turn may negatively influence the growth and development of the keystone species.

### Conclusion

Our results suggest that the fate of interspecies competition may depend on the size or age of the fish species (Munday et al., 2001; Byström and García-Berthou, 1999), and expressions of different behaviors (freeze or aggressive behaviors) may vary depending on the competitive ability of the interacting partner (Persson, 1983; Grether et al., 2009). Overall, the survival of a species greatly depends on the nature of its interactions within a given environment, and thus, exotic invasions to an ecosystem may not always be ecologically or environmentally threatening but also may impede the behavioral interactions of the natives within the introduced system.

## CRediT authorship contribution statement

**Suman Mallick:** Conceptualization, Methodology, Investigation, Formal analysis, Visualisation, Writing – original draft, Writing – review & editing. **Ratna Ghosal:** Conceptualization, Methodology, Visualisation, Writing – review & editing. **Jitendra Kumar Sundaray:** Methodology, Writing – review & editing.

## Data availability

All the data is available as tables and figures within the manuscript.

## Acknowledgments

We are thankful to the Scheme of Developing high quality research (SHODH), Department of Education, Government of Gujarat, India for providing fellowship to Suman Mallick.

---

**Supplementary Fig. 1.**
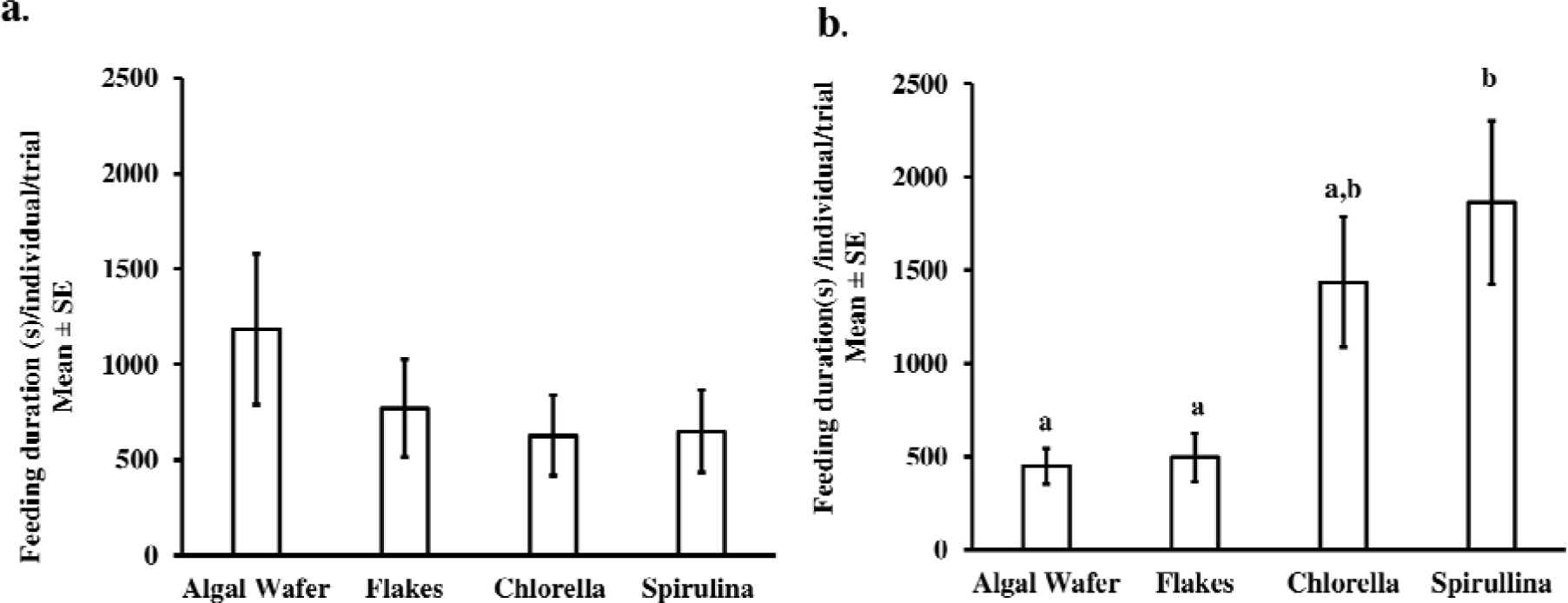
Feeding durations (Mean±SE per individual per trial, N=9 trials for each food item) of large-size Amazon sailfin catfish (panel a) in C trials and large-size rohu (panel b) in R trials for four different food items. Different alphabets represent significant difference (P≤0.05). ‘+’ represents the mean value per individual per trial.

**Supplementary Fig. 2.**
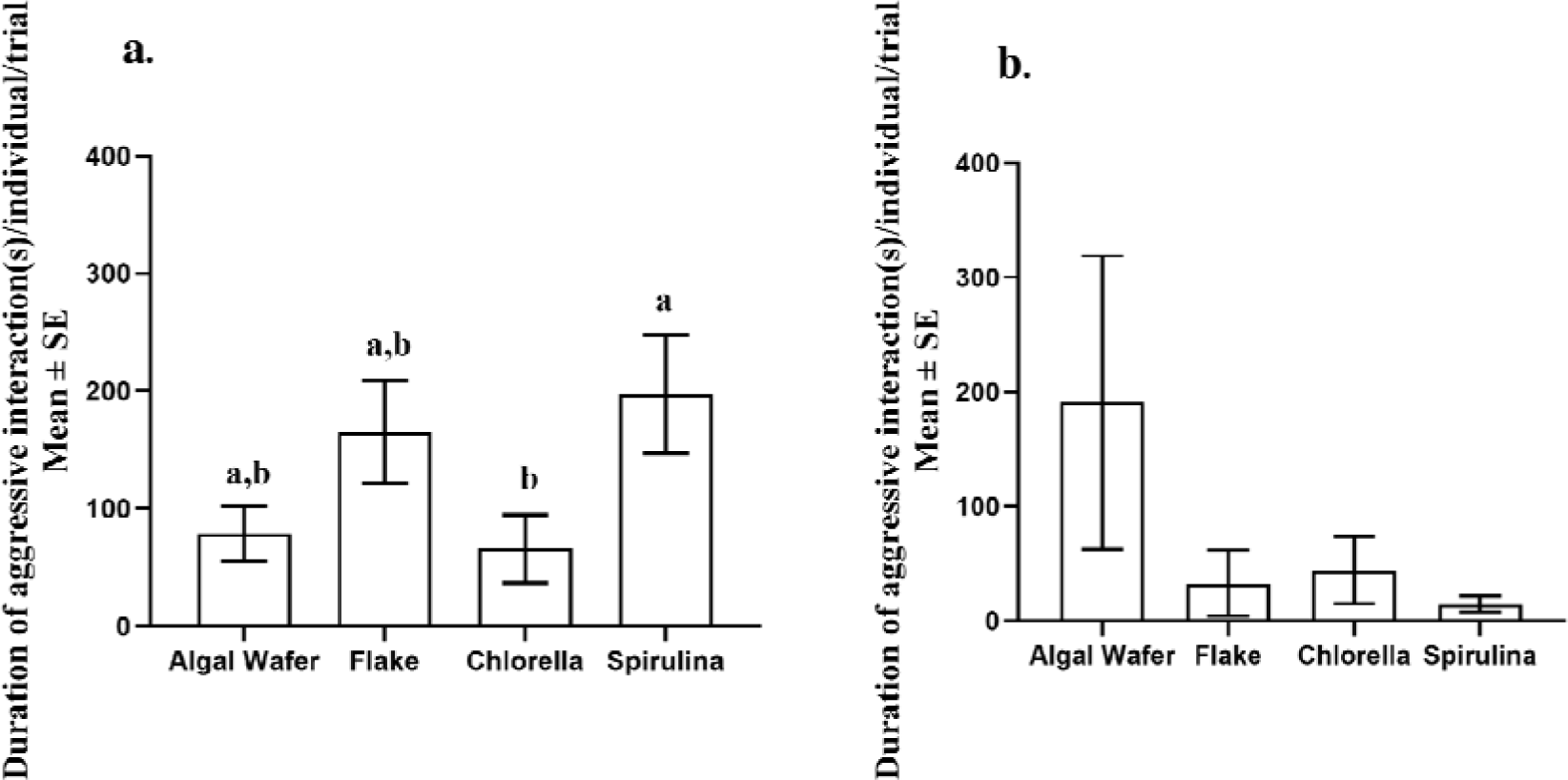
Duration of aggressive interactions (Mean±SE per individual per trial, N=9 trials for each food item) of large-size Amazon sailfin catfish (panel a) in C trials and large-size rohu (panel b) in R trials for four different food items. Different alphabets represent significant differences (P≤0.05). ‘+’ represents the mean value per individual per trial.

## References

Balshine, S., Verma, A., Chant, V. and Theysmeyer, T., 2005. Competitive interactions between round gobies and logperch. J. Great Lakes Res. 31(1), 68–77. 10.1016/S0380-1330(05)70238-0.

Bašić, T., Copp, G.H., Edmonds-Brown, V.R., Keskin, E., Davison, P.I., Britton, J.R., 2019. Trophic consequences of an invasive, small-bodied non-native fish, sunbleak *Leucaspius delineatus*, for native pond fishes. Biol. Invasions 21, 261–275. 10.1007/s10530-018-1824-y.

Bass, S.L., Gerlai, R., 2008. Zebrafish (*Danio rerio*) responds differentially to stimulus fish: the effects of sympatric and allopatric predators and harmless fish. Behav. Brain Res. 186(1), 107–117. 10.1016/j.bbr.2007.07.037.

Beisner, B.E., Ives, A.R., Carpenter, S.R., 2003. The effects of an exotic fish invasion on the prey communities of two lakes. J. Anim. Ecol. 72(2), 331–342. 10.1046/j.1365-2656.2003.00699.x.

Bhaumik, U., Mukhopadhyay, M.K., Shrivastava, N.P., Sharma, A.P., Singh, S.N., 2017. A case study of the Narmada River system in India with particular reference to the impact of dams on its ecology and fisheries. Aquat Ecosyst Health Manag 20(1-2), 151–159. 10.1080/14634988.2017.1288529.

Bijukumar, A., Smrithy, R., Sureshkumar, U., George, S., 2015. Invasion of South American suckermouth armoured catfishes *Pterygoplichthys* spp.(Loricariidae) in Kerala, India-a case study. J. Threat. Taxa 7(3), 6987–6995. 10.11609/JoTT.o4133.6987-95.

Birch, L.C., 1957. The meanings of competition. Am. Nat. 91(856), 5–18.

Borcherding, J., Heubel, K., Storm, S., 2019. Competition fluctuates across years and seasons in a 6-species-fish community: empirical evidence from the field. Rev. Fish Biol. Fish. 29, 589–604. 10.1007/s11160-019-09567-x.

Boujard, T., 1995. Diel rhythms of feeding activity in the European catfish, *Silurus glanis*. Physiol. Behav. 58(4), 641–645. 10.1016/0031-9384(95)00109-V.

Bowers, M.A., Brown, J.H., 1982. Body Size and Coexistence in Desert Rodents: Chance or Community Structure? Ecological Archives E063-002. Ecology 63(2), 391–400. 10.2307/1938957.

Brenden, T.O., Murphy, B.R., 2004. Experimental assessment of age-0 largemouth bass and juvenile bluegill competition in a small impoundment in Virginia. N. Am. J. Fish. Manag. 24(3),1058–1070. 10.1577/M03-115.1.

Brown, G.E., Godin, J.G.J., Pedersen, J., 1999. Fin-flicking behavior: a visual antipredator alarm signal in a characin fish, *Hemigrammus erythrozonus*. Anim. Behav. 58(3), 469–475. 10.1006/anbe.1999.1173.

Byström, P., García-Berthou, E., 1999. Density dependent growth and size specific competitive interactions in young fish. Oikos 86(2), 217–232. 10.2307/3546440.

Carrete, M., Lambertucci, S.A., Speziale, K., Ceballos, O., Travaini, A., Delibes, M., Hiraldo, F., Donázar, J.A., 2010. Winners and losers in humanLmade habitats: interspecific competition outcomes in two Neotropical vultures. Anim. Conserv. 13(4), 390–398. 10.1111/j.1469-1795.2010.00352.x.

Chaichana, R., Pouangcharean, S., Yoonphand, R., 2011. Habitat, abundance and diet of invasive suckermouth armored catfish (Loricariidae *Pterygoplichthys*) in the Nong Yai Canal, East Thailand. Trop. Zool. 24(1), 49–62.

Das, B.K., Ray, A., Manna, R.K., Roshith, C.M., Baitha, R., Karna, S.K., Gupta, S.D., Bhor, M., 2020. Occurrence of exotic vermiculated sailfin catfish *Pterygoplichthys disjunctivus* from the lower stretch of River Ganga, West Bengal, India. Curr. Sci. 119(12), 2006–2009. 10.18520/cs/v119/i12/2006-2009.

Das, M.K., Sharma, A.P., Vass, K.K., Tyagi, R.K., Suresh, V.R., Naskar, M., Akolkar, A.B., 2013. Fish diversity, community structure and ecological integrity of the tropical River Ganges, India. Aquat Ecosyst Health Manag 16(4), 395–407. 10.1080/14634988.2013.851592.

de Lourdes Ruiz-Gomez, M., Kittilsen, S., Höglund, E., Huntingford, F.A., Sørensen, C., Pottinger, T.G., Bakken, M., Winberg, S., Korzan, W.J., Øverli, Ø., 2008. Behavioral plasticity in rainbow trout (*Oncorhynchus mykiss*) with divergent coping styles: when doves become hawks. Horm Behav. 54(4), 534–538. 10.1016/j.yhbeh.2008.05.005.

Dwivedi, A.C., Mayank, P., Misra, V.K., Prakash, S., Mishra, A.S., 2020. Study on age and growth of Indian Major carp (*Labeo rohita*) from the Ganga River. J. krishi vigyan. 9(si), 276–279. 10.5958/2349-4433.2020.00118.X.

Dwivedi, A.C., Nautiyal, P., 2012. Stock assessment of fish species *Labeo rohita*, *Tor tor* and *Labeo calbasu* in the rivers of Vindhyan region, India. J. Environ. Biol. 33(2), 261–264.

Edwards, R.J., 2001. New additions and persistence of the introduced fishes of the upper San Antonio River, Bexar County, Texas. Tex. J. Sci. 53(1), 3–13.

Franco, M. and Arce, E., 2022. Aggressive interactions and consistency of dominance hierarchies of the native and nonnative cichlid fishes of the Balsas basin. Aggress. Behav. 48(1), 103–110. 10.1002/ab.21997.

Fullerton, A.H., Lamberti, G.A., Lodge, D.M. and Goetz, F.W., 2000. Potential for resource competition between Eurasian ruffe and yellow perch: growth and RNA responses in laboratory experiments. Trans. Am. Fish. Soc. 129(6), 1331–1339. 10.1577/1548-8659(2000)129%3C1331:PFRCBE%3E2.0.CO;2.

Gause, G. F., 1934. The struggle for existence. Williams and Wilkins, Baltimore. 163 p.

Gestring, K.B., Shafland, P.L., Stanford, M.S., 2010. Status of the exotic Orinoco sailfin catfish (*Pterygoplichthys multiradiatus*) in Florida. Fla. Sci. 73(2), 122–137.

Glova, G.J., 1986. Interaction for food and space between experimental populations of juvenile coho salmon (*Oncorhynchus kisutch*) and coastal cutthroat trout (*Salmo clarki*) in a laboratory stream. Hydrobiologia, 131(2), 155–168. 10.1007/BF00006779.

Grabowska, J., Kakareko, T., Błońska, D., Przybylski, M., Kobak, J. and Copp, G.H., 2016. Interspecific competition for a shelter between non-native racer goby and native European bullhead under experimental conditions–Effects of season, fish size and light conditions. Limnol. 56, 30–38.10.1016/j.limno.2015.11.004.

Grether, G.F., Losin, N., Anderson, C.N. and Okamoto, K., 2009. The role of interspecific interference competition in character displacement and the evolution of competitor recognition. Biol. Rev. 84(4), 617–635.10.1111/j.1469-185X.2009.00089.x

Herath, H.M.G.K., Kalutharage, N.K., Cumaranatunga, P.R.T., 2020. Solutions to an alien species invasion from aquarium aquaculture: Isolation and characterization of acid soluble collagen from sailfin catfish, *Pterygoplichthys disjuctivus* (Weber, 1991) in Sri Lanka. Sri Lanka. J. Aquat. Sci. 25(1), 19–32. 10.4038/sljas.v25i1.7573.

Hossain, M.Y., Vadas Jr, R.L., Ruiz-Carus, R., Galib, S.M., 2018. Amazon sailfin catfish *Pterygoplichthys pardalis* (Loricariidae) in Bangladesh: a critical review of its invasive threat to native and endemic aquatic species. Fishes 3(1), 1–12. 10.3390fishes3010014.

Hussan, A., Sundaray, J.K., Mandal, R.N., Hoque, F., Das, A., Chakrabarti, P.P., Adhikari, S., 2019. Invasion of non-indigenous suckermouth armoured catfish of the genus *Pterygoplichthys* (Loricariidae) in the East Kolkata Wetlands: stakeholders’ perception. Indian J. Fish. 66(2), 29–42. 10.21077/ijf.2019.66.2.86267-05.

Ichinose, K., 2002. Influence of age and body size on alarm responses in a freshwater snail *Pomacea canaliculata*. J. Chem. Ecol. 28, 2017–2028. 10.1023/A:1020749911877

Janssen, A.B., Teurlincx, S., An, S., Janse, J.H., Paerl, H.W., Mooij, W.M., 2014. Alternative stable states in large shallow lakes?. J. Great Lakes Res. 40(4), 813–826. 10.1016/j.jglr.2014.09.019.

Jumawan, J.C., Vallejo, B.M., Herrera, A.A., Buerano, C.C., Fontanilla, I.K.C., 2011. DNA barcodes of the suckermouth sailfin catfish *Pterygoplichthys* (Siluriformes: Loricariidae) in the Marikina River system, Philippines: Molecular perspective of an invasive alien fish species. Philipp. Sci. Lett. 4(2), 103–113.

Levin, B.A., Phuong, P.H., Pavlov, D.S., 2008. Short communication Discovery of the Amazon sailfin catfish *Pterygoplichthys pardalis* (Castelnau, 1855)(Teleostei: Loricariidae) in Vietnam. J. Appl. Ichthyol. 24, 715–717. 10.1111/j.1439-0426.2008.01185.x.

Ludlow, M.E., Walsh, S.J., 1991. Occurrence of a South American armored catfish in the Hillsborough River, Florida. Fla. Sci. 54(1), 48–50.

Mahamood, M., Javed, M., Alhewairini, S.S., Zahir, F., Sah, A.K., Ahmad, M.I., 2021. *Labeo rohita*, a bioindicator for water quality and associated biomarkers of heavy metal toxicity. NPJ Clean Water 4(1), 1–7.

Mallick, S., Hussan, A., Sundaray, J.K., Ghosal, R., 2023. Ecological assessment of the Amazon sailfin catfish (*Pterygoplichthys species*) within the Indian freshwaters: a mesocosm-based approach.bioRxiv 10.1101/2023.07.22.550140.

McCarthy, I. D., D. J. Gair, D. F. Houlihan. Feeding rank and dominance in Tilapia *Coptodon rendalli* under defensible and indefensible patterns of food distribution. J. Fish Biol. 55(4), 854–867. 10.1016/j.anbehav.2013.10.002.

Mills, M.D., Rader, R.B. and Belk, M.C., 2004. Complex interactions between native and invasive fish: the simultaneous effects of multiple negative interactions. Oecologia 141, 713–721.10.1007/s00442-004-1695-z.

Miyai, C.A., Sanches, F.H.C., Costa, T.M., Colpo, K.D., Volpato, G.L. and Barreto, R.E., 2011. The correlation between subordinate fish eye colour and received attacks: a negative social feedback mechanism for the reduction of aggression during the formation of dominance hierarchies. Zool. 114(6), 335–339. 10.1016/j.zool.2011.07.001

Modeel, S., Joshi, B.D., Yadav, S., Bharti, M., Negi, R.K., 2023. Mitochondrial DNA reveals shallow population genetic structure in economically important Cyprinid fish *Labeo rohita* (Hamilton, 1822) from South and Southeast Asia. Mol. Biol. Rep. 50(6), 4759–4767. 10.1007/s11033-023-08442-0.

Mohanta, K.N., Subramanian, S., Komarpant, N., Saurabh, S., 2008. Alternate carp species for diversification in freshwater aquaculture in India. Aquac. Asia 13(1), 11–15.

Mookerji, N., Rao, T.R., 1993. Patterns of prey selection in rohu (*Labeo rohita*) and singhi (*Heteropneustes fossilis*) larvae under light and dark conditions. Aquaculture 118(1-2), 85–104. 10.1016/0044-8486(93)90283-5.

Munday, P.L., Jones, G.P., Caley, M.J., 2001. Interspecific competition and coexistence in a guild of coralLdwelling fishes. Ecol. 82(8), 2177–2189. 10.1890/0012-9658(2001)082[2177:ICACIA]2.0.CO;2.

Munilkumar, S., Nandeesha, M.C., 2007. Aquaculture practices in Northeast India: Current status and future directions. Fish Physiol. Biochem. 33, 399–412. 10.1007/s10695-007-9163-4.

Näslund, J., Pettersson, L., Johnsson, J.I., 2016. Behavioural reactions of three-spined sticklebacks to simulated risk of predation—effects of predator distance and movement. Facets 1(1), 55–66. 10.1139/facets-2015-0015.

Ngasotter, S., Panda, S.P., Mohanty, U., Akter, S., Mukherjee, S., Waikhom, D., Devi, L.S., 2020. Current scenario of fisheries and aquaculture in India with special reference to Odisha: a review on its status, issues and prospects for sustainable development. Int. j. bio-resour. stress manag.11(4), 370–380. 10.23910/1.2020.2126a.

Nol, E., Smith, J.N., 1987. Effects of age and breeding experience on seasonal reproductive success in the song sparrow. J. Anim. Ecol. 56(1), 301–313. 10.2307/4816

Parvez, M.T., Lucas, M.C., Hossain, M.I., Chaki, N., Mohsin, A.B.M., Sun, J., Galib, S.M., 2023. Invasive vermiculated sailfin catfish (*Pterygoplichthys disjunctivus*) has an impact on highly valued native fish species. Biol. Invasions 25, 1795–1809. 10.1007/s10530-023-03012-8.

Paszkowski, C.A., 1985. The foraging behavior of the central mudminnow and yellow perch: the influence of foraging site, intraspecific and interspecific competition. Oecologia 66, 271–279. 10.1007/BF00379865.

Persson, L., 1983. Effects of intra-and interspecific competition on dynamics and size structure of a perch (*Perca fluviatilis)* and a roach (*Rutilus rutilus)* population. Oikos 41(1) 126–132. 10.2307/3544354.

Persson, L., Greenberg, L.A., 1990. Juvenile competitive bottlenecks: the perch (*Perca fluviatilis)*Lroach (*Rutilus rutilus*) interaction. Ecol. 71(1), 44–56.10.2307/1940246

Piria, M., Simonović, P., Kalogianni, E., Vardakas, L., Koutsikos, N., Zanella, D., Ristovska, M., Apostolou, A., Adrović, A., Mrdak, D., Tarkan, A.S., 2018. Alien freshwater fish species in the Balkans—Vectors and pathways of introduction. Fish Fish 19(1), 138–169. 10.1111/faf.12242.

Qasim, A.M., Jawad, L.A., 2022. Presence of the Amazon sailfin catfish, *Pterygoplichthys pardalis* (Castelnau, 1855)(Pisces: Loricariidae), in the Shatt al-Arab River, Basrah, Iraq. Integrative Systematics: Stuttgart Contributions to Natural History, 5(1), 95–103. 10.18476/2022.647187.

Rabbane, M.G., Kabir, M.A., Habibullah-Al-Mamun, M. and Mustafa, M.G., 2022. Toxic effects of arsenic in commercially important fish Rohu Carp, *Labeo rohita* of Bangladesh. Fishes 7(5), 2–17. 10.3390/fishes7050217.

Rahman, M.M., 2015. Effects of co-cultured common carp on nutrients and food web dynamics in rohu aquaculture ponds. Aquac. Environ. Interact. 6(3), 223–232. 10.3354/aei00127.

Rahman, M.M., Verdegem, M.C., Nagelkerke, L.A., Wahab, M.A., Verreth, J.A., 2008. Swimming, grazing and social behaviour of rohu *Labeo rohita* (Hamilton) and common carp *Cyprinus carpio* (L.) in tanks under fed and non-fed conditions. Appl. Anim. Behav. Sci. 113(1-3), 255–264. 10.1016/j.applanim.2007.09.008.

Rao, K.R., Sunchu, V., 2017. A report on *Pterygoplichthys pardalis* Amazon sailfin suckermouth Catfishes in freshwater tanks at Telangana state, India. Int. j. fish. aquat. sci. 5(2), 249–254.

Raphael, S., Aude, J., Olivier, P., Mélissa, P.L., Aurélien, B., Christophe, B., Jean Marc, P., Sylvie, B., Alain, D., Wilfried, S., 2016. Characterization of a genotoxicity biomarker in three-spined stickleback (*Gasterosteus aculeatus* L.): Biotic variability and integration in a battery of biomarkers for environmental monitoring. Environ. Toxicol. 31(4), 415–426. 10.1002/tox.22055.

Rasal, K.D., Sundaray, J.K., 2020. Status of genetic and genomic approaches for delineating biological information and improving aquaculture production of farmed rohu, *Labeo rohita* (Ham, 1822). Rev. Aquac. 12(4), 2466–2480. 10.1111/raq.12444.

Rudolf, V.H., McCrory, S., 2018. Resource limitation alters effects of phenological shifts on inter-specific competition. Oecologia 188(2), 515–523. 10.1007/s00442-018-4214-3.

Saba, A.O., Rasli, N.F., Ismail, A., Zulkifli, S.Z., Ghani, I.F.A., Muhammad-Rasul, A.H., Azmai Amal, M.N., 2020. A Report on Introduced Amazon Sailfin Catfish, *Pterygoplichthys pardalis* in Gombak Basin, Selangor, with Notes on Two Body Patterns of the Species. Pertanika J. Trop. Agric. Sci. 43(4), 693–703.

Šárová, R., Špinka, M., Stěhulová, I., Ceacero, F., Šimečková, M., Kotrba, R., 2013. Pay respect to the elders: age, more than body mass, determines dominance in female beef cattle. Anim. Behav. 86(6), 1315–1323. 10.1016/j.anbehav.2013.10.002

Savino, J.F., Kolar, C.S., 1996. Competition between nonindigenous ruffe and native yellow perch in laboratory studies. Trans. Am. Fish. Soc. 125(4), 562–571. 10.1577/1548-8659(1996)125%3C0562:CBNRAN%3E2.3.CO;2.

Seshagiri, B., Swain, S.K., Pillai, B.R., Satyavati, C., Sravanti, Y., Rangacharyulu, P.V., Rathod, R., Ratnaprakash, V., 2021. Suckermouth armoured catfish (*Pterygoplichthys* spp.) menace in freshwater aquaculture and natural aquatic systems in Andhra Pradesh, India. Int. J. Fish. Aquat. 9(1), 375–384. 10.22271/fish.2021.v9.i1e.2423.

Seshagiri, B., Kumar, A., Pradhan, P.K., Sood, N., Kumar, U., Satyavati, C., Sravanti, Y., Prasoon, K., Ghosh, A., Kantharajan, G., Basheer, V.S., 2022. Farming practices and farmers’ perspective of a non-native fish red-bellied Pacu, *Piaractus brachypomus* (Cuvier,1818) in India. Aquaculture 547, 737483. 10.1016/j.aquaculture.2021.737483.

Singh, A.K., Lakra, W.S., 2011. Ecological impacts of exotic fish species in India. Aquac. Asia 16(2), 23–25.

Sumanasinghe, H.W., Amarasinghe, U.S., 2014. Population dynamics of accidentally introduced Amazon sailfin catfish, *Pterygoplichthys pardalis* (Siluriformes, Loricariidae) in Pologolla reservoir, Sri Lanka.Sri Lanka J. Aquat. Sci. 18, 37–45.

Suresh, V.R., Ekka, A., Biswas, D.K., Sahu, S.K., Yousuf, A., Das, S., 2019. Vermiculated sailfin catfish, *Pterygoplichthys disjunctivus* (Actinopterygii: Siluriformes: Loricariidae): invasion, biology, and initial impacts in east Kolkata Wetlands, India. Acta Ichthyol. Piscat. 49(3), 221–233. 10.3750/AIEP/02551.

Tarkan, A.S., Vilizzi, L., Top, N., Ekmekçi, F.G., Stebbing, P.D., Copp, G.H., 2017. Identification of potentially invasive freshwater fishes, including translocated species, in Turkey using the Aquatic Species Invasiveness Screening Kit (ASLISK). Int. Rev. Hydrobiol. 102(1-2), 47–56. 10.1002/iroh.201601877.

Templeton, C.N., Greene, E. and Davis, K., 2005. Allometry of alarm calls: black-capped chickadees encode information about predator size. Science 308(5730), 1934–1937. 10.1126/science.1108841

Umar, M.T., Ramayani, S., 2021. Population dynamics of sailfin catfish (*Pterygoplichthys sp.* Hancock, 1828) in sidenreng lake water, Sidenreng Rappang District, South Sulawesi. In IOP Conference Series: Earth and Environmental Science 860(1), 1–8. 10.1088/1755-1315/860/1/012104.

Wakida-Kusunoki, A.T., Ruiz-Carus, R., Amador-del-Angel, E., 2007. Amazon sailfin catfish, *Pterygoplichthys pardalis* (Castelnau, 1855)(Loricariidae), another exotic species established in southeastern Mexico. Southwest. Nat. 52(1), 141–144.

Winemiller, K.O., Taylor, D.H., 1987. Predatory behavior and competition among laboratory-housed largemouth and smallmouth bass. Am. Midl. Nat. 117(1),148–166. 10.2307/2425716.

